# Species composition and environmental adaptation of indigenous Chinese cattle

**DOI:** 10.1101/170449

**Authors:** Yahui Gao, Mathieu Gautier, Xiangdong Ding, Hao Zhang, Yachun Wang, Xi Wang, MD Omar Faruque, Junya Li, Shaohui Ye, Xiao Gou, Jianlin Han, Johannes A. Lenstra, Yi Zhang

## Abstract

Indigenous Chinese cattle combine taurine and indicine origins and occupy a broad range of different environments. By 50K SNP genotyping we found a discontinuous distribution of taurine and indicine cattle ancestries with extremes of less than 10% indicine cattle in the north and more than 90% in the far south and southwest China. Model-based clustering and *f4*-statistics indicate introgression of both banteng and gayal into southern Chinese cattle while the sporadic yak influence in cattle in or near Tibetan area validate earlier findings of mitochondrial DNA analysis. Geographic patterns of taurine and indicine mitochondrial and Y-chromosomal DNA diversity largely agree with the autosomal cline. The geographic distribution of the genomic admixture of different bovine species is proposed to be the combined effect of prehistoric immigrations, gene flow, major rivers acting as genetic barriers, local breeding objectives and environmental adaptation. Whole-genome scan for genetic differentiation and association analyses with both environmental and morphological covariables are remarkably consistent with previous studies and identify a number of genes implicated in adaptation, which include *TNFRSF19*, *RFX4*, *SP4* and several coat color genes. We propose indigenous Chinese cattle as a unique and informative resource for gene-level studies of climate adaptation in mammals.

## Introduction

China harbors around 10 million of indigenous cattle^1^. It is commonly referred as yellow cattle and divided into 53 indigenous breeds raised in various agro-ecological environments^2,3.^ Their diversity and unique species composition emerged from a complex history. Domestic cattle spread to East Asia by at least two routes. Taurine cattle migrated from north Eurasia to northern China and northeast Asia between 5000 and 4000 BP^4^. This is supported by the evidence that ancient cattle from northern China, dated 4500 to 2300 BP, carried only taurine mtDNA haplotypes^5^. A unique mtDNA haplotype, T4, observed in East Asian cattle breeds^6-9^ is a subtype of the common haplogroup T3^10^, suggesting a founder effect in Chinese taurine cattle^11^.

Indicine cattle (zebu) migrated eastward from their domestication center in the Indus valley and entered China from the south since 3000 BP^4,12^. Southeast Asian and southern Chinese cattle are morphologically and genetically recognized as zebu^13,14^. Yue *et al.* provided evidence for an additional southwestern immigration route of zebu from India into northwest China^15^.

The taurine and indicine cattle migrations resulted in a morphological gradient from humpless taurine cattle in the north to humped indicine cattle in southern and southwestern China^3^. This has been confirmed by genetic studies using mtDNA^7,8,16,17^ and Y-linked markers^18^. A genetic diversity study using microsatellite markers clustered Chinese indigenous cattle breeds into one taurine and four indicine groups^19^.

In addition to taurine and indicine cattle, several other bovine species have been living in southern China and Southeast Asia, including banteng, gaur or gayal, which may have been the dominant cattle species until 4500 BP^4,12^. Gayal in Yunnan province of China carried indicine or taurine mtDNA but gaur Y chromosome, indicating its hybrid origin^20^. Meanwhile, in Tibetan Autonomous Region (TAR) of China, bidirectional introgression between yak and cattle has also been reported^21-24^. Genetic admixture has been identified between zebu and Bali cattle (domestic banteng) in Indonesia^22,25^. A previous study on hair color and blood protein polymorphism provided evidence of banteng introgression into Hainan cattle in southeastern China^26^, which was confirmed by genomic SNP array data^25^.

Genomic SNP array has become a powerful tool for population genomics studies in animals. A recent genomic variation study by Decker *et al.* revealed a worldwide pattern of genetic admixture in domestic cattle^25^. Other studies focused on Creole^27^, American^28^, East African zebu^29^ and Korean cattle^30^. These advanced approaches also allow the genomic localization of genes involved in the adaptation to natural or artificial selective constraints^27,31-33^. In the current study, we generated 50K SNP genotypes to infer the fine-scale characterization of unique species composition of highly diverse Chinese cattle. In addition, we performed a whole genome-scan for adaptive differentiation and association analyses with environmental and morphological population-specific covariables to detect genes that responded to adaptive constraints.

## Results

### Genomic variation

Observed heterozygosity (Table 1) ranged from 0.145 to 0.327 in Chinese cattle populations (Fig. 1). Mongolian cattle (MG, NM) and Hazak cattle had the highest values but southern and southwestern Chinese cattle populations were the lowest. This is most likely explained by the ascertainment bias, by which the heterozygosity of indicine cattle is underestimated. Indeed, the observed heterozygosity correlates negatively (*r^2^*=0.96) with the zebu ancestry. A similar trend has previously been observed in West-African cattle^34^. Bali cattle (0.026), gayal (0.059) and yak (0.029) also have relatively low levels of heterozygosity as normally observed with SNP panels designed for a different species^35^.

**Table 1.**
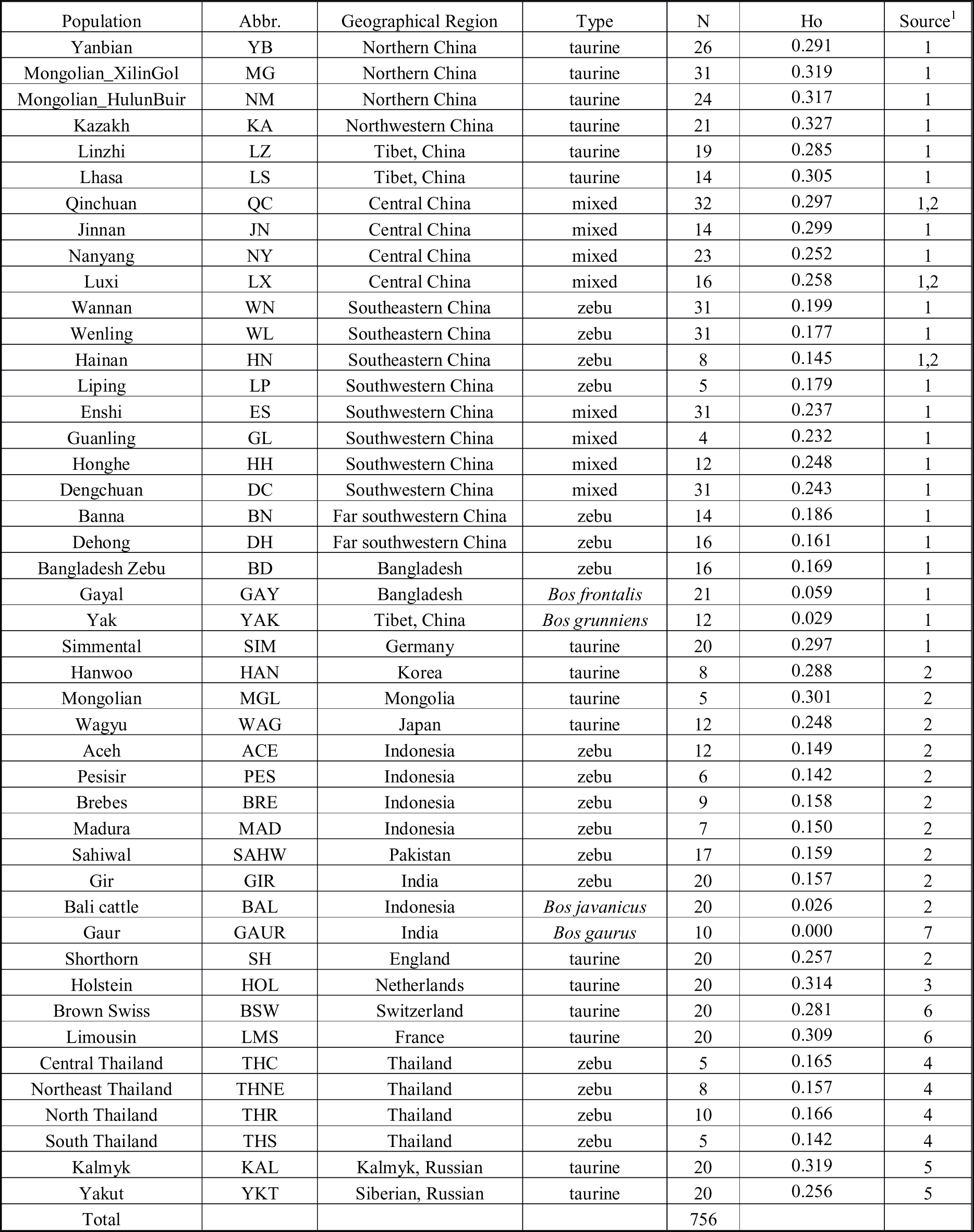
Sampling information of different cattle populations and their observed heterozygosity (Ho). Source of data. 1) this study; 2) Decker *et al*. (2014); 3) Gautier *et al*. (2010); 4) Wangkunmhang *et al*. (2015); 5) Decker *et al*. (2016); 6) Matukumalli *et al*. (2009); 7) Decker *et al*. (2009).

**Figure 1.**
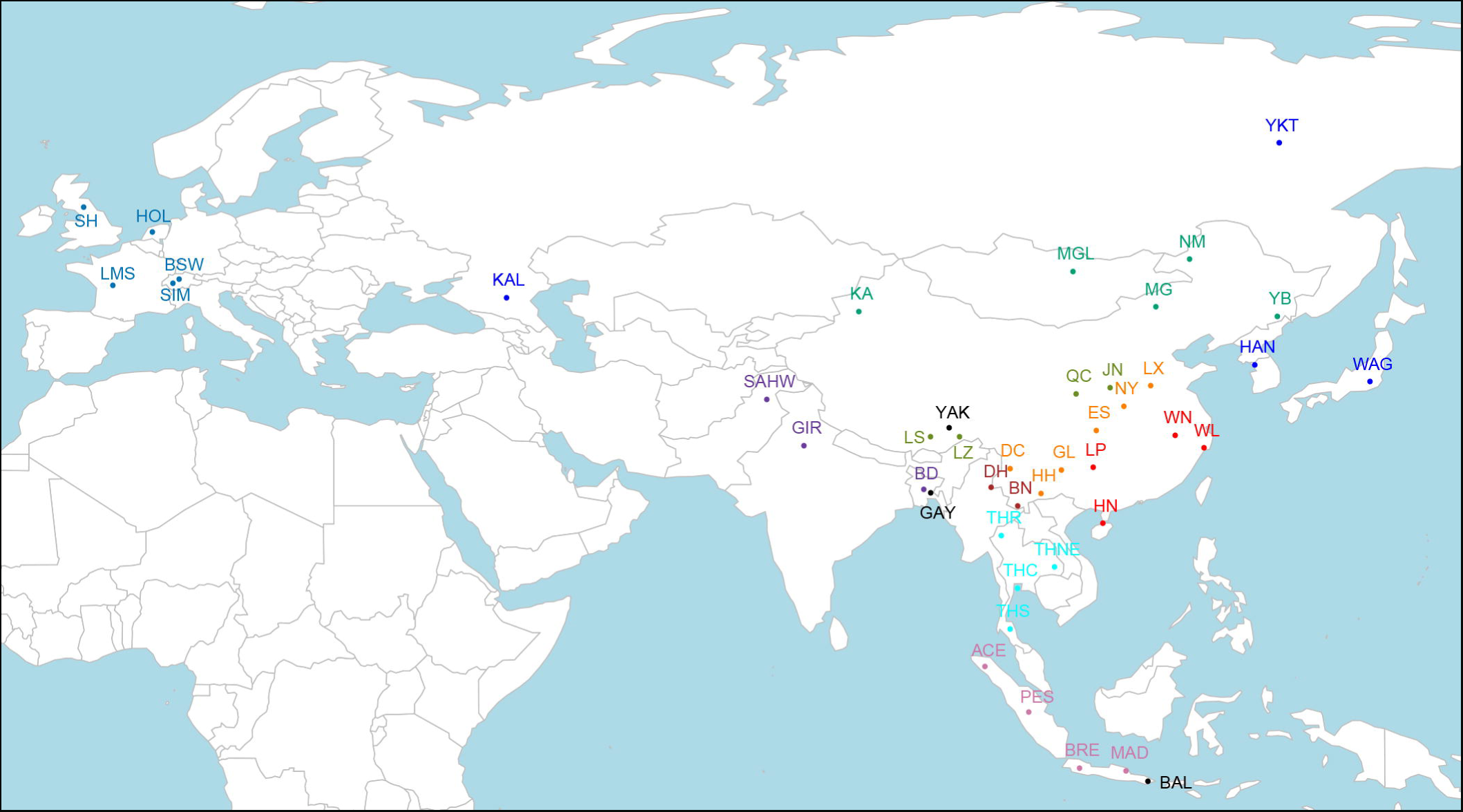
Geographical distribution of cattle populations. Detailed information is showed in Table 1. Map was created using R 3.3.1 packages *rworldmap*, *maps* and *mapproj* (https://cran.rproject.org/web/packages/). Package *rworldmap* was used to generate outline and colorful dots while packages *maps* and *mapproj* were used to add texts.

### Population structure

Five methods were implemented in this part to explore the population structure.

#### PCA

Figure 2a shows a scatter plot of the first two principal components (PCs), allowing to assess the structuring of genetic diversity across all the 45 sampled populations, including outgroup species (Bali cattle, yak, gaur and gayal). The first PC accounting for 17.1% of total variation separates taurine and indicine cattle as well as other bovine species. The predominant taurine breeds from northern China and TAR cluster with European breeds while populations from southeastern and far-southwestern China are closer to Indian zebu. The second component accounting for 3.1% of all variation displays the contrast of Chinese cattle to other bovine species (Bali, gayal and yak) and also differentiates European cattle breeds. The intermediate positions of LZ, MAD, BRE, Bali and gayal populations indicate gene flow between species.

**Figure 2.**
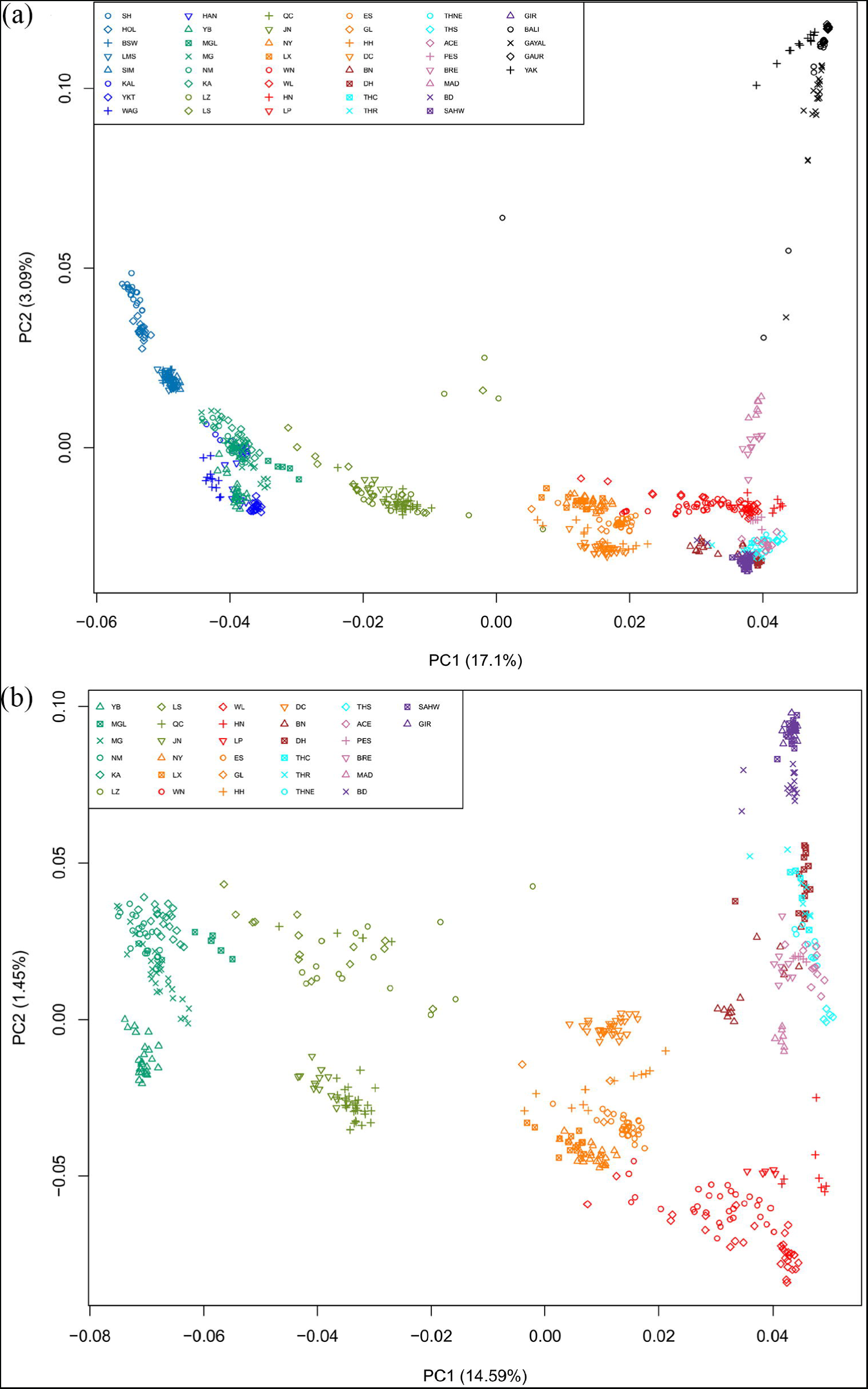
PCA plots describing the relationships among populations. (a) Whole data set of all the 45 sampled populations. (b) A subset of data including only Asian cattle.

In an analysis of only Asian cattle (Fig. 2b), the first PC that explains 14.5% of the genetic variation again corresponds to the taurine-indicine separation, but the second PC that explains 1.45% of the genetic variation represents a gradient from India via Southeast Asia to Southeastern China.

#### Model-based clustering

Figure 3a shows an unsupervised hierarchical clustering performed on the whole data set (i.e., including outgroup species) with different values of K, the number of clusters. K=2 reproduces the first PCA coordinate with taurine and indicine clusters and a taurineindicine gradient from north to south. Higher values of K differentiate European from Asian taurine breeds (K=3), zebu from other bovine species (K=4), and southern Chinese from Asian indicine cattle (K=5). Increasing K further generates separate clusters for European cattle breeds whereas Chinese cattle populations tend to show admixed ancestries. The cross-validation error gave the lowest value at K = 17 (Supplementary Fig. S1), which differentiates Bali cattle, gayal, yak, the European taurine breeds and indicine cattle from India/Pakistan, Southeast Asia and southeastern China, respectively.

**Figure 3.**
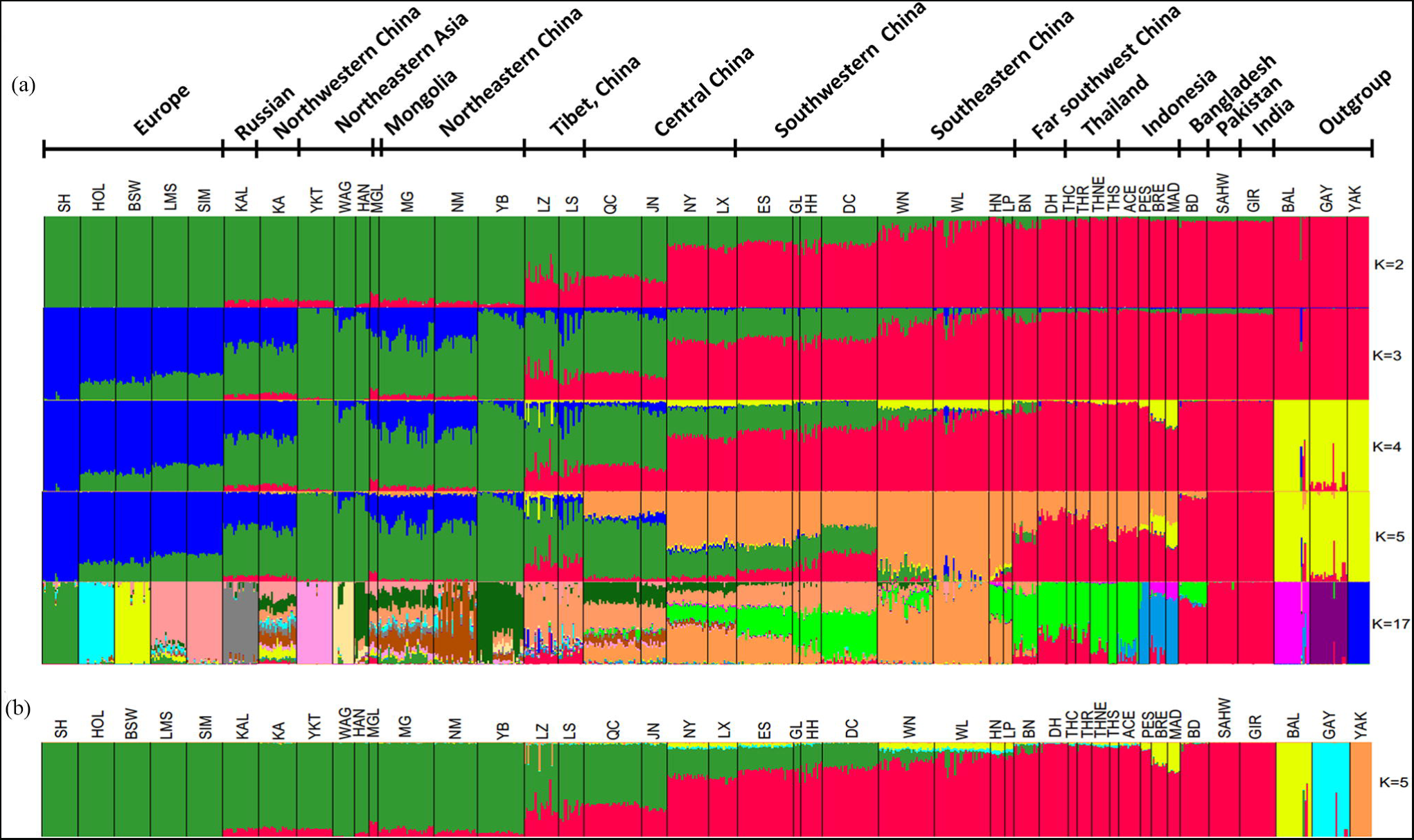
Ancestries and population structuring of Chinese cattle revealed by (a) unsupervised Admixture analysis (K=2,3,4,5,17) and (b) supervised Admixture analysis (K=5). For supervised admixture analysis, five European cattle breeds (SH, HOL, LMS, SIM) were set to represent taurine ancestry whereas GIR and SAHW represented indicine ancestry. BAL, GAY and YAK were three outgroup bovine species. The putative hybrid animals in Bali and Gayal detected by unsupervised admixture analysis were excluded from the pure ancestry.

Figure 3b shows a supervised clustering with prior population information for taurine breeds (SH, HOL, BSW, SIM), indicine breeds (GIR, SAHW) and other bovine species. In this analysis, Bali cattle represents the banteng and the domestic gayal replaces the wild gaur since the latter was too inbred for this analysis. This analysis reproduces the zebu-banteng hybridization in Indonesia^36^ and yak introgression into Tibetan cattle, but also suggests minor banteng and gayal and southeastern China (Supplementary Fig. S2). Supplementary Figure S3 shows the geographic distribution of the inferred species components.

#### NeighborNet network

In a NeighborNet graph constructed from the matrix of Reynolds’ distances between populations using Splitstree (Fig. 4), European and Indian cattle are at the extreme ends of the network, which is entirely in agreement with both the first PCA coordinate and the K=2 clustering. Figure 4 also reproduces the intermediate positions of the predominantly taurine-indicine breeds from TAR or central China and also of the predominantly indicine breeds from central or southwestern China. In addition, the network confirms the affinity of Indonesian and southeastern Asian continental breeds with Bali cattle and gayal.

**Figure 4.**
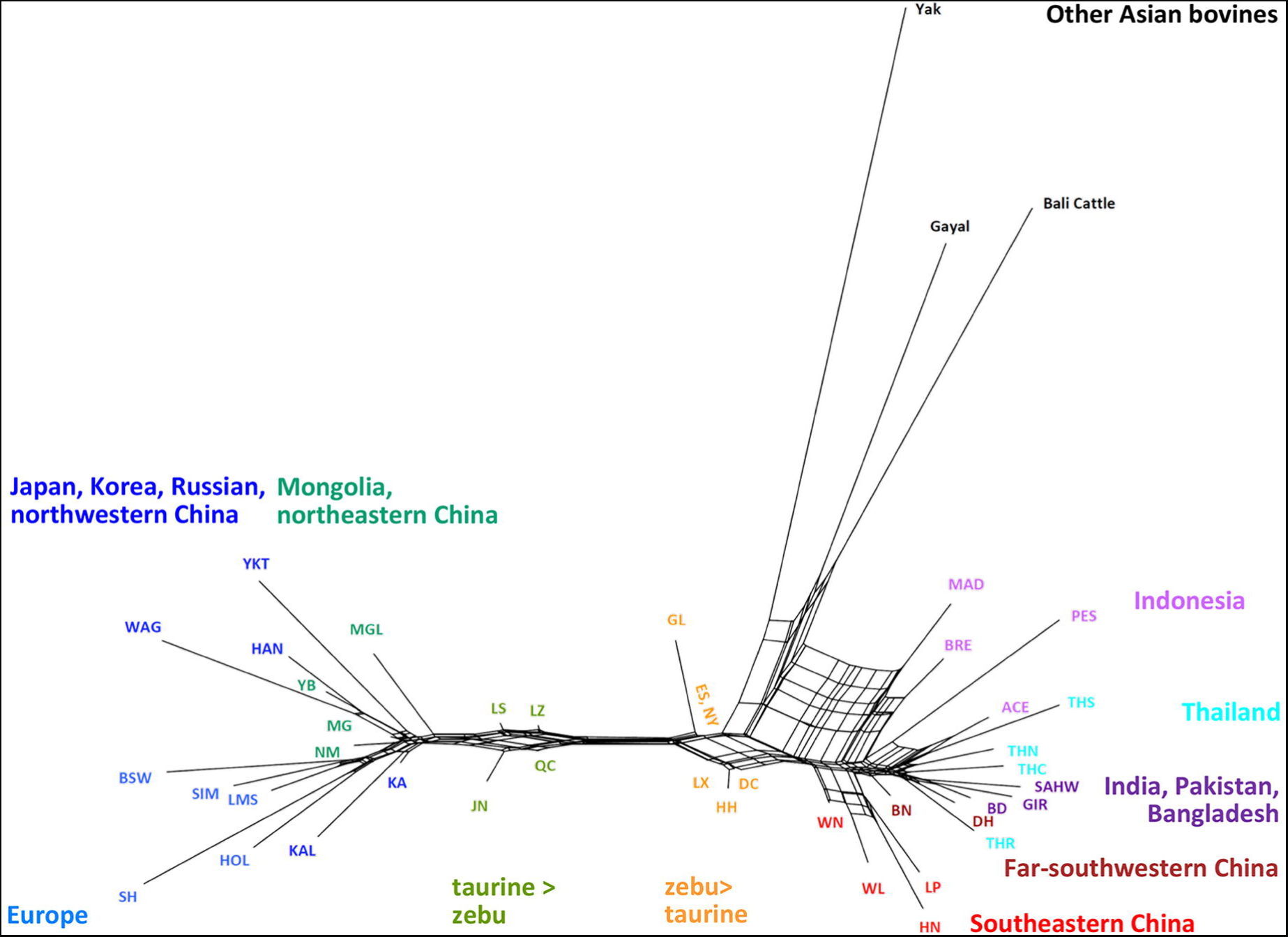
NeighborNet graph of 44 cattle populations. An allele frequency-dependent distance metric (Reynolds) was used to construct the NeighborNet.

#### Population mixture

The taurine-indicine mixed composition of Chinese cattle was confirmed by the sensitive *f4* ratio test (Supplementary Table S1). The estimated taurine ancestry ranges from close to 0.0 in southeastern China to close to 1.0 in north, with intermediate values ranging from 0.282 to 0.760, from 0.240 to 0.320 and from 0.656 to 0.770 in central China, southwestern China and TAR, respectively. This pattern was highly correlated with the average memberships estimated by model-based clustering (*r*=998, Supplementary Table S1).

As shown in Fig. 5a, negative values of the four-population *f4*-statistics (GIR, X; BAL,YAK) suggests gene flow from Bali to indicine populations in southeastern China and Southeast Asia. This gene flow has clearly been more consequential for the Indonesian cattle breeds, MAD, BRE and PES as well as the southern Chinese breeds LP, HN, WN and WL. For the Indonesian breeds this confirms previous results of Mohamad *et al.*^36^ and Decker *et al.*^25^. Although Bali cattle are relatively closely related to gaur, replacing Bali cattle by gaur as source of admixture generates only a moderately negative value for the Indonesian breeds (Fig. 5b). This is even observed for Bali cattle as a test breed and may reflect the inbreeding of the gaur samples. However, the same plot shows relatively low (*i.e.*, negative, indicating gene flow) values for the southern Chinese breeds. In combination with the supervised model-based clustering (Fig. 3b), this may suggest that these breeds have been introgressed by gaur and/or gayal in addition to banteng, the wild ancestor of Bali cattle.

**Figure 5.**
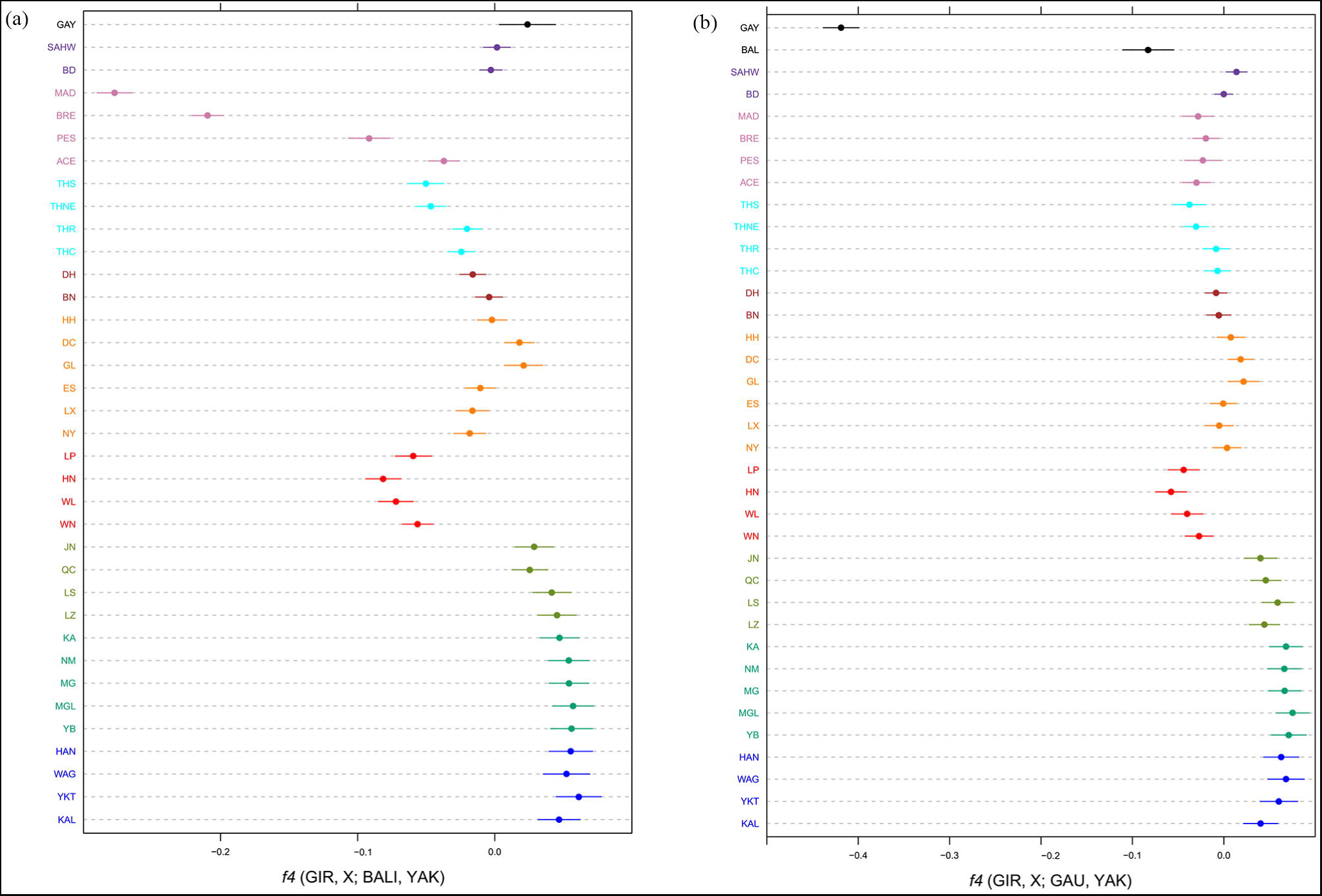
Visualization of *f4*-statistics. (a) *f4*-statistics of the form D (GIR, X; BAL, YAK) identified gene flow from Bali to other breeds. (b) *f4*-statistics of the form D (GIR, X; GAU, YAK) identified gene flow from Gaur to other breeds. The whiskers represent the standard error.

A *f4*(SH,X;GAY,YAK) plot (Supplementary Fig. S4A) generated negative values for the indicine GIR, SAHW and BD as test breeds. Since this is not observed with the wild gaur instead of the domestic gayal (Supplementary Fig. S4B), this indicates indicine introgression into the domestic gayal population. This is confirmed by two other observations:

- The statistic *f4*(GAU,GAY;X,YAK) is negative for all test breeds, but clearly more negative for indicine than for taurine breeds (Supplementary Fig. S4C).
- *f4*(GIR,BAL;GAY,YAK) is positive but *f4*(GIR,BAL;GAU,YAK) is not (Supplementary Fig. S4D; Fig. 5B). The same patterns are also observed if GIR is replaced by SH (Supplementary Fig. S4A, B), which is phylogenetically close to the indicine GIR. Apparently the allele sharing of GAY with GIR or SH outweighs the allele sharing expected because of the phylogenetic relationship of gayal and Bali cattle, which most likely is an effect of the ascertainment bias of the SNP panel towards taurine breeds^37^.

#### Uniparental markers

Supplementary Table S2 and Figure S5 show mtDNA and Y-chromosomal haplotype distributions of Chinese cattle breeds, based on our data supplemented by literature data (Supplementary Table S3). These uniparental markers show a north-to-south taurine-indicine gradient that resembles closely the autosomal cline (Fig. 6). Haplotype diversity (Supplementary Table S2 and Fig. S5C) shows that the diversity of taurine mtDNA hardly decreased from north to south, while indicine mtDNA clearly decreases from southeastern and southwestern China to northern China.

**Figure 6.**
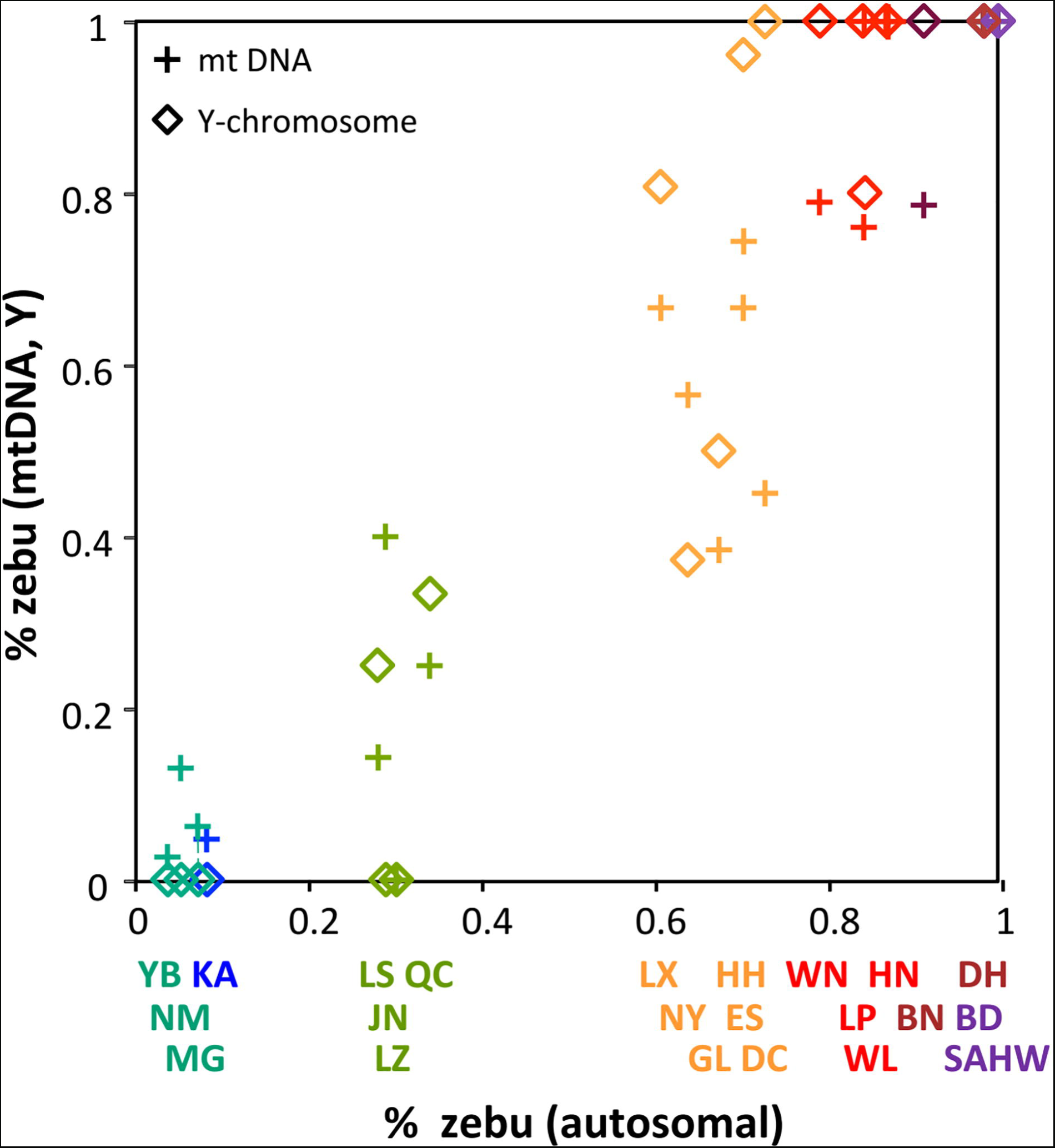
Correlation of autosomal, mtDNA and Y-chromosomal indicine components. Uniparental markers were used to build the correlation graph which showed a north-to-south taurine-indicine gradient that resembles closely the autosomal cline.

### Detection of genomic regions subjected to adaptive constraints

For a genome-wide scan for adaptive differentiation, the XtX differentiation statistics was estimated for each SNP under the so-called core model and visualized in a Manhattan plot (Fig. 7). Among 84 significant SNPs, SNP Hapmap28985-BTA-73836 on BTA5 was the most significant.

**Figure 7.**
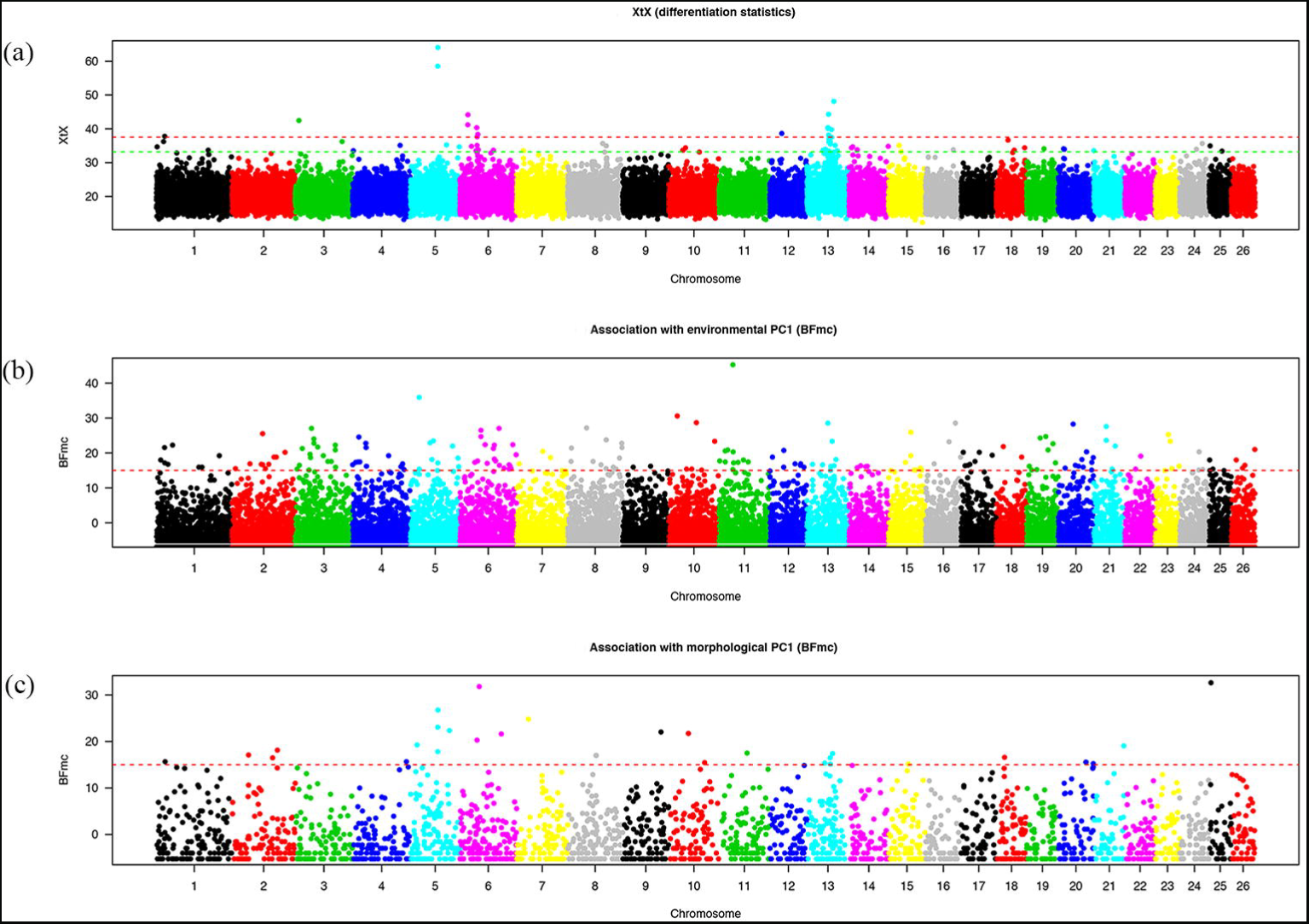
Whole genome scan for adaptive divergence and association analyses in 20 Chinese cattle breeds. (a) Manhattan plot of the XtX statistics. (b) Manhattan plot of the BFmc (association with environmental PC1). (c) Manhattan plot based on the BFmc (association with morphological PC1).

The three synthetic environmental covariables are associated with 185 significant SNPs (Supplementary Table S4), from which SNP Hapmap44345-BTA-119580 on BTA11 was the most significant (Fig. 7). From 30 SNPs associated with morphological covariables (Supplementary Table S4), SNP ARS-BFGL-NGS-67505 on BTA25 was most significant (Fig. 7).

As a matter of expedience, we applied a sliding window approach to identify the main genomic regions of interest as described in Gautier^38^. Briefly, the UMD3.1 bovine genome assembly^39^ on which all the SNPs were mapped was first split into 4718 consecutive 1-Mb windows (with a 500-kb overlap). For each window, we counted the number n_s_ of SNPs that were either significantly differentiated at the 0.1% threshold (i.e., with XtX > 32.3) or associated (BF >15) with at least one of the four population-specific covariables. Windows with n_s_>=2 were deemed significant and overlapping windows were further merged. This yielded 27 significant regions detailed in Table 2, of which eight showed significantly differentiated SNPs; 22, 0 and 4 displayed SNPs associated with the first, the second and the third environmental co-variable, respectively; and 4 displayed SNPs associated with the morphological covariable. Note that only 3 regions (out of the 8) containing significantly differentiated SNPs did not contain any SNPs associated with the population-specific covariables studied. Finally, a total of 28 candidate genes were annotated in the significant regions using UCSC (https://genome.ucsc.edu/) (Table 2).

**Table 2.**
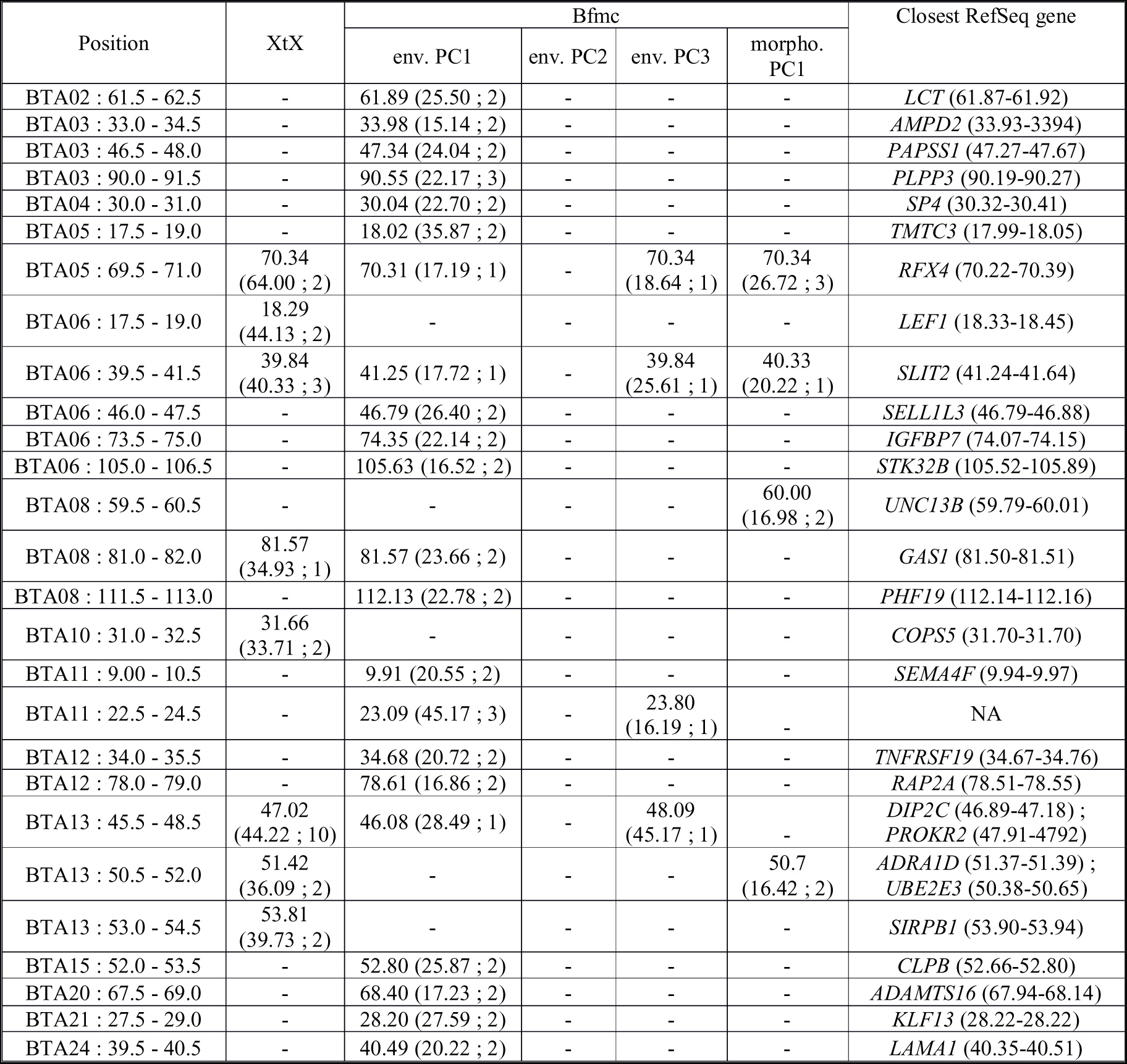
Regions consisting of overlapping windows each containing at least one SNP with XtX score >32.2 or at least one Bfmc score >15 identified by BayPass whole genome scan. For each XtX or Bfmc test, the table gives the peak position in Mb as well as the peak statistics value and the number of SNPs in parentheses with a test value above the corresponding threshold. Dash (–) indicates non-significant results.

## Discussion

We have investigated the species composition and genomic, mitochondrial as well as Ychromosomal variation in Chinese cattle populations. A clear north-south gradient of taurine and indicine cattle ancestries combined with banteng, gayal and yak introgressions into southern and southwestern Chinese cattle populations defines the pattern of admixture among Chinese indigenous cattle, which are supposed to underlie local breeding objectives and adaptation to different agro-ecological environments. Genome scan for adaptive differentiation and association with population-specific covariable identify regions and candidate genes relevant for environmental adaption and morphological differentiation in Chinese cattle.

Previous genetic diversity studies using mtDNA and Y-linked markers have characterized a north-south gradient of taurine and indicine admixture in Chinese cattle, which is consistent with the transition from humpless to humped morphology^7,8,16-18^. A microsatellite study differentiated five groups of Chinese indigenous cattle breeds^19^.

Our PCA (Fig. 2) and model-based clustering (Fig. 3, Supplementary Fig. S3) patterns as well as the NeighborNet graph (Fig. 4) reveal a clear transition from taurine cattle in the north to zebu in the south with consistent admixture levels within the breeds and a clear demarcation of five groups:

- Cattle from Manchuria, Inner Mongolia and northwestern China with 4 to 8% indicine admixture. This group corresponds to the taurine type 1^19^ (this type also comprises Tibetan cattle) and to Group 9B in the Felius classification^40^.
- Taurindicine cattle in TAR and northern China above the Yellow or Wei River with 28 to 35% indicine admixture, denoted as zebu type 2^19^ and belonging to the Huanghuai group of central Chinese cattle (Group 10A)^40^.
- Taurindicine cattle below the Yellow River with 61 to 73% indicine genome, corresponding to indicine types 1 and 4^19^. The northernmost LX, JX and NY breeds belong to the Huanghai group (Group 10A)^40^, but the other breeds to the Changzu group (Group 10B)^40^.
- Predominant indicine cattle in southern China with an indicine genomic component of 79 to 87% (zebu type 3^19^, Group 10B^40^).
- Indicine cattle with a >90% indicine genome (also indicine type 3^19^, Group 10B^40^).

Cattle from Northeast Asia and northern China have typical taurine morphological features. Its genetic distinctiveness and old origin evolved from the unique mtDNA haplogroup T4 found in modern breeds from Japan, Korea, Mongolia^6,41^, Siberia (Yakut cattle)^9^， northern China^7,8^ (Supplementary Fig. S5A) and in ancient cattle in northern China from 2300-4500 BP^5^. In addition, Decker *et al*. found a separate position of Japanese and Korean cattle^25^. In this study, we found that northern Chinese, South Korean and Japanese cattle share genetic ancestry with the Siberian Yakut and the indigenous northeastern China close to Korea. These breeds represent the eastern range of Turano-Mongolian cattle, which have retained their original dark-brown coat color pattern in Mongolian and Korean cattle.

The admixture pattern, however, suggests European cattle influence to northern Chinese cattle diversity. Possibly migrations of nomads in the steppes of Central Asia and Mongolia, which in the Middle Ages led to the establishment of the Mongolian empire, facilitated eastward as well as westward gene flow across Eurasia. Additionally, in the past few decades programs have been implemented in China to upgrade productivity by crossing local breeds with European breeds^3,12^. Influence of European cattle was captured by model-based clustering analysis, e.g. Brown Swiss and Simmental in Kazak, Holstein in LS, and Limousine in LX, JN and WL (Fig. 3).

From north to south, levels of taurine autosomal, mitochondrial and Y-chromosomal DNA decrease, but are still appreciable in southwestern China (Supplementary Figs S3 and S5). The occurrence of indicine mtDNA in mixed taurine-indicine cattle is a unique feature of Chinese breeds^11^. The high taurine mtDNA diversity in southern China (Supplementary Fig. S5C) indicates an absence of a major founder effect. This is more compatible with immigration before the arrival of indicine cattle around 3000 BP than with a later introgression into an existing indicine population. The immigration of zebus from the present Myanmar and Indochina to the north with a plausible contribution of an eastward gene flow in western China^15^ resulted in significant indicine components in northern Chinese and Mongolian cattle (Supplementary Figs S3 and S5). The pattern of indicine mtDNA diversity (Supplementary Fig. S5C) suggests a population bottleneck when they crossed the Pearl River in southeastern China.

Y-chromosomal and autosomal indicine components correlate well (Fig. 6). Exceptions are the Guanlin (GL) and Nanyang (NY) breeds with relatively low indicine Y-chromosomal frequencies. Remarkably, the indicine autosomal component has a discontinuous distribution (Figs 2-4; Supplementary Fig. S3) with the largest gap across the Yellow River separating JN (29% zebu) from ES (61%) and NY (64%). This river might be a physical barrier of gene flow, but this cannot explain the absence in our panel of cattle with an indicine component between 35% and 61%. It might be hypothesized that in cattle with equal taurine and indicine components, outbreeding depression outweighs heterosis, for instance by incompatibility of genes from different origins conferring fitness^42^. It is again remarkable that cattle at both sides of the river are categorized as belonging to the Huanghuai group and resemble western Asian and African cattle because of their similar cervico-thoracic hump^2^.

The NeighborNet shows that indicine cattle from far southwestern China (DH, BN), which are found in the west of the Mekong River, are more closely related to Thai cattle than to southeastern Chinese cattle. This is also supported by the PCA and admixture patterns (Figs 2 and 3). The separate position of Thai cattle confirms the results of Wangkunmhang *et al*.^14^. The Mekong River acted also as a genetic barrier for swamp buffalo^43^.

Another potential source of diversity in southern Chinese cattle are the introgressions from other bovine species living in China and Southeast Asia, including yak, gayal and its wild ancestor gaur, and banteng, represented by its domestic relative Bali cattle. (Fig. 3; Supplementary Fig. S2). There are several examples of hybridization of different bovine species. Yak mtDNA has been detected in indigenous cattle distributed on the Qinghai-Tibetan plateau and in Diqing cattle (DQ) of Yunnan province^21,23^. We did not detect yak mtDNA in our cattle panel, but we detected influence of yak in the Tibetan LZ population based on the model-based clustering analysis.

A study of blood protein polymorphism^26^ suggested banteng ancestry in Hainan cattle. Using mtDNA and microsatellite genotyping, Mohamad *et al*.^44^ characterized banteng admixture in Indonesian zebu breeds. This was confirmed by 50K SNP analysis^25^, which also detected a low level of banteng introgression in southern Chinese breeds. However, analyzing Chinese cattle together with both banteng (represented by its domestic derivative Bali cattle) and gaur (the wild ancestor of gayal) with model based clustering (Fig. 3) and *f4*-statistics (Fig. 5) provided consistent evidence of both gayal and banteng introgression into WL, WN and HN breeds from southeastern China and also into Thai zebu at a relatively low proportion.

Conversely, similar *f4*-statistics (Supplementary Fig. S4) suggested introgression of zebu into gayal, which has been confirmed in the gayal population from Yunnan carrying both indicine and taurine mitochondrial genomes^20^. Similarly, cattle introgression has been detected in yak populations^24^.

The genetic variation described above reflects the combined effect of prehistoric immigrations of taurine and indicine cattle, subsequent gene flow between populations, local selection objectives and environmental adaptation. Indigenous Chinese cattle with indicine-taurine ratios varying between zero and one and subject to a broad range of climates is a valuable resource to identify potential genomic regions and functional genes underlying the environmental adaption. By combining signals of population differentiation (XtX) and association with three synthetic environmental covariables and one synthetic morphological covariable (Supplementary Fig. S6), we identified 27 genomic regions and 28 candidate genes targeted by natural or artificial selection (Table 2).

Interestingly, 12 out of the 27 regions overlap with core selective sweep (CSS) regions^45^, while 20 and 23 regions overlap with breed-wise and breed group-wise hotspots of selective sweeps, respectively^46^ (Supplementary Table S5). However, we report for the first time the region BTA12: 34.0-35.5 Mb, which harbors *TNFRSF19*. This member of the TNF-receptor superfamily^47^ is highly expressed during embryonic development^48^.

In other studies, strong signals of selection in tropical cattle have been detected on BTA5^32,34,49,50^. Notably, Porto-Neto *et al.* identified a 20 Mb region on BTA5 with effects on parasite resistance, yearling weight, body condition score, coat color and penile sheath score^32^. We found a significant signature of selection for XtX and the environmenmtal PC1 and PC3 as well as the morphological PC1 in the region 69.5-71.0 on BTA5 region 69.5-71.0 (Table 2), which contains the candidate gene *RFX4*. This gene is a member of Regulatory Factor X (RFX) family of transcriptional regulators that influence MHC class II expression^51^ and play a critical role in brain development^19,52^. It was also found to affect heifer fertility in tropical composition breed Brangus^53^.

Coat color is an important target of selection in many domestic animals. The common denotation of yellow cattle for the indigenous Chinese cattle refers to its predominant light to dark brown color. In current study, selection signatures were identified near several known color genes, including *KITLG* (near SNP BTA-74300-no-rs on BTA5)^38,54^, and *LEF1*^32,55^ (here indicated by a peak in the XtX GWAS on BTA6). These genes and another candidate gene *MCM6* (near ARS-BFGL-NGS-92772 on BTA2, also identified by Hudson *et al.*^50^) overlap with pigmentation QTL regions underlying UV-protection^56^. The environmental PC1 signal near *IGFBP7* and the combined XtX-morphological signal near *ADRA1D* (Table 2) are close to the coat color genes *KIT*^56,57^ and *ATRN*^56,58^, respectively.

We further detected an environmental PC1 association signal near *SP4* (Sp4 transcription factor) as novel candidate gene on BTA4. It is a member of the Sp1-family of zinc finger transcription factors and is required for normal murine growth, viability, and male fertility^59^. In cattle, *SP4* was suggested to have effect on body size and testicular growth from birth to yearling age^60^.

It is interesting to note that Chinese and African cattle have developed independently a variable taurine-indicine ancestry following a gradient from tropical to temperate climates. An attractive opportunity is a detailed comparison of gene variants involved in climate adaptation by using whole genome sequence data^61^. It may be anticipated that in both regions adaptation to agroecological constraints is mediated by recruiting and combining gene variants from taurine and indicine origins with possible original contributions in Chinese indigenous cattle from the indicine mtDNA, and the minor gayal, and banteng and yak genomic ancestry.

## Methods

### Ethics statement

The protocols for collection of the blood and hair samples of experimental individuals were reviewed and approved by the Institutional Animal Care and Use Committee (IACUC) at China Agricultural University. And all experiments were performed in accordance with approved relevant guidelines and regulations.

### Samples collection and genotyping

We collected samples of 437 animals from 24 breeds (Table 1), twenty of which are indigenous cattle populations from northeastern China, central China, southeastern China, southwestern China, far southwestern China and or TAR (Fig. 1). We also examined Bangladeshi cattle and German Simmental, and two related bovine species, the gayal and yak. Samples were genotyped with Illumina BovineSNP50 BeadChip using standard procedures^62^. Genotypes are accessible via the WIDDE repository (http://widde.toulouse.inra.fr/widde/).

We compared these newly generated data with published genotypes of some European and Asian cattle breeds, Bali cattle and gaur (Table 1). The combined data set comprises 37,429 SNPs. Using PLINK^63^, we removed SNPs with call rates <90% or with minor allele frequencies <0.001 and discarded individuals with 10% missing genotypes. The resulting data set contained 36,872 SNPs and 736 animals from 44 populations representing taurine cattle, zebu, and three species related to cattle (*Bos javanicus* - banteng, *Bos gruniens* - yak and, *Bos frontalis* - gayal). Gaur (*Bos gaurus*)^64^ was used instead of gayal in *f4* analysis (see below) of the species composition of Chinese cattle because of the indicine zebu introgression into gayal.

### Population genetic analysis

We used PLINK^63^ to calculate the observed homozygosity for each population. Three complementary methods were used to analyze the genetic diversity among populations. First, a Principal Component Analysis (PCA) was carried out to investigate the pattern of genetic differentiation among populations and individuals using the R package SNPRelate^65^, which performs eigen-decomposition of the genetic covariance matrix to compute the eigenvalues and eigenvectors. Second, population structure was evaluated by unsupervised and supervised model-based hierarchical clustering implemented in the Admixture software^66^. The results were visualized using the program Distruct^67^. Third, a NeighbourNet network was constructed using Reynold’s distances between populations using Splitstree 4.13^68^.

To investigate species composition of Chinese cattle, we used the four-population test (*f4-*statistics) implemented in ADMIXTOOLS^69^. Additionally, taurine and indicine ancestries in Chinese cattle populations were quantified via the *f4* ratio estimation in ADMIXTOOLS, which allows inference of the admixture proportions without access to accurate surrogates for the ancestral populations^69^. The proportion of taurine ancestry was then computed as

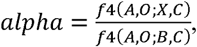

in which O is an outgroup (BAL), B a reference taurine cattle (YKT), C an Indian zebu cattle (GIR), A a population related to B (SH), and X the Chinese target population. Standard errors were computed with the Block Jackknife procedure in ADMIXTOOLS using default options^69^.

### Mitochondrial DNA and Y-chromosomal markers

A 445-bp mtDNA control region was amplified and sequenced as described previously (GenBank accession codes KY682307-KY682687)^20^. These sequences were analyzed together with published sequences (Supplementary Tables S2 and S3). Haplotype diversity of the segment 16,023-16,262 (numbering according to GenBank accession no. V00654) was computed using the software DnaSP^70^. Y-chromosomal genotyping was carried out with the protocol described by Bonfiglio *et al*. (2012)^71^, which differentiates Y1 (dominant in north Europe) and Y2 (dominant in other taurine cattle) and Y3 (indicine cattle) type Y chromosome^72^.

### Genome-scan for adaptive differentiation and association with environmental and morphological covariables

Whole genome-scans for adaptive differentiation and association with population-specific co-variables were performed with BayPass 2.1^38^. The underlying models explicitly account for the covariance structure among the population allele frequencies, which make the approach particularly robust to complex demographic histories^38^. Identification of overly differentiated SNPs was based on the XtX statistics^38,73^ estimated under the core model of BayPass. To calibrate the XtX’s, a pseudo-observed data set (POD) containing 250,000 SNPs simulated under the inference model with hyperparameters equal to those estimated on the real data set was generated and further analyzed under the same conditions following the procedure described in Gautier^38^. In particular, we ensured that the posterior estimate of the scaled covariance matrix of population allele frequencies (Omega) obtained with the POD was similar to that obtained on the real data since the FMD distance between the two matrices was found equal to 0.28^38^. Similarly, the posterior means of the two hyperparameters a and b for the *Beta* distribution of across population allele frequencies obtained on the POD (a=b=1.02) were almost equal to the ones obtained in the original data (a=b=1.00). Taken together, these sanity checks indicated that the POD faithfully mimics the real data set, allowing us to define a 0.1% significance threshold on the XtX statistics (XtX=32.3) to identify genomic regions harboring footprints of selection.

Genome-wide analysis of association with population specific co-variables was carried out using the default options of the AUX model parameterized with the scaled covariance matrix (Omega) obtained on the real data set as described above. This model allows to account explicitly for multiple testing issue by integrating over (and estimating) the unknown proportion of SNPs actually associated with a given covariable. The support for association of each SNP with each covariable was evaluated by computing Bayes Factor (BF) and a BF>15 is considered as decisive evidence for association^38^.

We collected values for six environmental covariables, i.e. average temperature, average relative humidity, sunshine, average air pressure, wind speed, and precipitation from China Meteorological Administration (http://data.cma.cn/). Values for 10 morphological covariables, i.e. male body weight, female body weight, male height, female height, male body length, female body length, male heart girth, female heart girth, male fore-shank circumference, and female fore– shank circumference were provided for 17 out of the 20 studied indigenous breeds in National Commission of Animal Genetics Resources^3^, but these data were not available for the breeds LZ, BN and HH. We carried out a PCA on scaled variables for environmental and morphological covariables separately. The first three environmental PCs and the first morphological PC were retained as uncorrelated co-variables for association studies (Supplementary Tables S6 and S7).

## Acknowledgement

This work was supported by National Natural Scientific Foundation of China (31561143010, 31272418) and China Agricultural Research System (CARS-37). Xi Wang was supported by Shanxi Scientific and Technological Development Programs (20140311018-1, 20120311021-1). The authors thank Dr. Hideyuki Mannen (Kobe University, Japan), Dr. Albano Beja-Pereira (Universidade do Porto, Portugal) and Hartati Hartati (Indonesian Agency for Agricultural Research and Development, Ministry of Agriculture) for providing detailed information on their data sets.

## Author Contributions

Y.Z., J.A.L. and M.G. conceived the experiments. Y.G., M.G., X.D., H.Z., Y.W., X.W., M.O.F., J.L., S.Y., X.G., J.H., J.A.L. and Y.Z. contributed samples. Y.G., M.G., J.A.L. and Y.Z. analysed the data. Y.G., M.G., J.A.L. and Y.Z. prepared the original draft. J.H., J.A.L. and Y.Z. reviewed and edited the paper.

## References

1. Zhang, Y. Calmly thinking on the cognition of the Chinese indigenous yellow cattle breed resources and sustainable utilization. China Cattle Science. 39, 1–6 (2013). In Chinese.

2. Chen, Y. C. & Cao, H. H. Diversity of Chinese yellow cattle breeds and their conservation. Biodiversity Science. 9, 275–283 (2001). In Chinese.

3. China National Commission of Animal Genetics Resources. Animal genetic resources in China – Bovine, (China Agriculture Press, 2011). In Chinese.

4. Payne, W. J. A. & Hodges, J. Tropical Cattle, Origins, Breeds and Breeding Policies, (Blackwell Science, 1997).

5. Cai, D. W., et al. The origins of Chinese domestic cattle as revealed by ancient DNA analysis. J. Archaeol. Sci. 41, 423–434 (2014).

6. Mannen, H., et al. Independent mitochondrial origin and historical genetic differentiation in North Eastern Asian cattle. Mol. Phylogenet. Evol. 32, 539–544 (2004).

7. Lei, C. Z., et al. Origin and phylogeographical structure of Chinese cattle. Anim. Genet. 37, 579–582 (2006).

8. Lai, S. J., Liu, Y. P., Liu, Y. X., Li, X. W. & Yao, Y. G. Genetic diversity and origin of Chinese cattle revealed by mtDNA D-loop sequence variation. Mol. Phylogenet. Evol. 38, 146–154 (2006).

9. Kantanen, J., et al. Maternal and paternal genealogy of Eurasian taurine cattle (*Bos taurus*). Heredity. 103, 404–415 (2009).

10. Achilli, A., et al. Mitochondrial genomes of extinct aurochs survive in domestic cattle. Curr. Biol. 18, R157–158 (2008).

11. Lenstra, J. A., et al. Meta-analysis of mitochondrial DNA reveals several population bottlenecks during worldwide migrations of cattle. Diversity. 6, 178–187 (2014).

12. Felius, M., Beerling, M. L., Buchanan, D. S., Theunissen, B., Koolmees, P. A. & Lenstra, J. A. On the history of cattle genetic resources. Diversity. 6, 705–750 (2014).

13. Chen, S., et al. Zebu cattle are an exclusive legacy of the South Asia. Mol. Biol. Evol. 27, 1–6 (2010).

14. Wangkumhang, P., et al. Genetic analysis of Thai cattle reveals a Southeast Asian indicine ancestry. PeerJ. 3, e1318, 10.7717/peerj.1318 (2015).

15. Yue, X., et al. When and how did *Bos indicus* introgress into Mongolian cattle? Gene. 537, 214–219 (2014).

16. Cai, X., Chen, H., Lei, C., Wang, S., Xue, K. & Zhang B. MtDNA diversity and genetic lineages of eighteen cattle breeds from *Bos taurus* and *Bos indicus* in China. Genetica. 131, 175–183 (2007).

17. Jia, S., et al. A new insight into cattle's maternal origin in six Asian countries. J Genet Genomics. 37, 173–180 (2010).

18. Li, R., et al. Paternal origins of Chinese cattle. Anim. Genet. 44, 446–469 (2013).

19. Zhang, G. X., et al. Genetic diversity and population structure of indigenous yellow cattle breeds of China using 30 microsatellite markers. Anim. Genet. 38, 550–559 (2007).

20. Gou, X., Wang, Y., Yang, S., Deng, W. & Mao H. Genetic diversity and origin of Gayal and cattle in Yunnan revealed by mtDNA control region and SRY gene sequence variation. J. Anim. Breed. Genet. 127, 154–160 (2010).

21. Yu, Y., Nie, L., He, Z. Q., Wen, J. K., Jian, C. S. & Zhang, Y. P. Mitochondrial DNA variation in cattle of South China: origin and introgression. Anim. Genet. 30, 245–250 (1999).

22. Kikkawa, Y., et al. Phylogenies using mtDNA and SRY provide evidence for male-mediated introgression in Asian domestic cattle. Anim. Genet. 34, 96–101 (2003).

23. Shi, J., Qiao, H. S., Hosoi, E. & Ozawa, S. Phylogenetic relationship between the Yellow cattle in Qinghai province, China, and Japanese Black cattle based on mitochondrial DNA D-loop sequence polymorphism. Anim. Sci. J. 75, 13–519 (2004). In Japanese.

24. Qi, X. B., Jianlin, H., Wang, G., Rege, J. E. O. & Hanotte, O. Assessment of cattle genetic introgression into domestic yak populations using mitochondrial and microsatellite DNA markers. Anim. Genet. 41, 242–252 (2009).

25. Decker, J. E., et al. Worldwide patterns of ancestry, divergence, and admixture in domesticated cattle. PLoS Genet. 10, e1004254 (2014).

26. Chen, Y., Wang, Y., Cao, H., Pang, Z. & Yang, G. Black-ear gene and blood polymorphism in four southern Chinese cattle groups. Anim. Genet. 25 Suppl 1, 89–90 (1994).

27. Gautier, M. & Naves, M. Footprints of selection in the ancestral admixture of a New World Creole cattle breed. Mol. Ecol. 20, 3128–3143 (2011).

28. McTavish, E. J., Decker, J. E., Schnabel, R. D., Taylor, J. F. & Hillis, D. M. New World cattle show ancestry from multiple independent domestication events. Proc. Natl. Acad. Sci. U.S.A. 110, E1398–1406 (2013).

29. Mbole-Kariuki, M. N., et al. Genome-wide analysis reveals the ancient and recent admixture history of East African Shorthorn Zebu from Western Kenya. Heredity. 113, 297–305 (2014).

30. Sharma, A., Lee, S. H., Lim, D., Chai, H. H., Choi, B. H. & Cho, Y. A genome-wide assessment of genetic diversity and population structure of Korean native cattle breeds. BMC Genet. 17, 139 (2016).

31. Ramey, H. R., Decker, J. E., McKay, S. D., Rolf, M. M., Schnabel, R. D. & Taylor, J.F. Detection of selective sweeps in cattle using genome-wide SNP data. BMC Genomics. 14, 382 (2013).

32. Porto-Neto, L. R., et al. The genetic architecture of climatic adaptation of tropical cattle. PLoS One. 9, e113284 (2014).

33. Flori, L., et al. Adaptive admixture in the West African bovine hybrid zone: insight from the Borgou population. Mol. Ecol. 23, 3241–3257 (2014).

34. Gautier, M., et al. A whole genome Bayesian scan for adaptive genetic divergence in West African cattle. BMC Genomics. 10, 550 (2009).

35. Miller, J. M., Kijas, J. W., Heaton, M. P., McEwan, J. C. & Coltman, D. W. Consistent divergence times and allele sharing measured from cross-species application of SNP chips developed for three domestic species. Mol Ecol Resour. 12, 1145–1150 (2012).

36. Mohamad, K., et al. The origin of Indonesian cattle and conservation genetics of the Bali cattle breed. Reprod. Domest. Anim. 47 Suppl 1, 18–20 (2012).

37. Lipson, M., & Reich, D. A working model of the deep relationships of diverse modern human genetic lineages outside of Africa. Mol. Biol. Evol. 34, 889–902 (2017).

38. Gautier, M. Genome-wide scan for adaptive divergence and association with population-specific covariates. Genetics. 201, 1555–1579 (2015).

39. Liu, Y., et al. Bos taurus genome assembly. BMC Genomics. 10, 180 (2009).

40. Felius, M., Koolmees, P. A. & Theunissen, B. On the breeds of cattle-historic and current classifications. Diversity. 2011;3(4):660–92.

41. Sasazaki, S., Odahara, S., Hiura, C., Mukai, F. & Mannen, H. Mitochondrial DNA variation and genetic relationships in Japanese and Korean cattle. Asian-Aust. J. Anim. Sci. 19, 1394–1398 (2006).

42. Todesco, M., et al. Hybridization and extinction. Evol Appl. 9, 892–908 (2016).

43. Zhang, Y., et al. Strong and stable geographic differentiation of swamp buffalo maternal and paternal lineages indicates domestication in the China/Indochina border region. Mol. Ecol. 25, 1530–1550 (2016).

44. Mohamad, K., et al. On the origin of Indonesian cattle. PLoS One. 4, e5490 (2009).

45. Gutiérrez-Gil, B., Arranz, J. J. & Wiener, P. An interpretive review of selective sweep studies in Bos taurus cattle populations: identification of unique and shared selection signals across breeds. Front Genet. 6, 167 (2015).

46. Randhawa, I. A., Khatkar, M. S., Thomson, P. C. & Raadsma, H. W. A meta-assembly of selection signatures in cattle. PLoS One. 11, e0153013 (2016).

47. Hu, S., Tamada, K., Ni, J., Vincenz, C. & Chen, L. Characterization of TNFRSF19, a novel member of the tumor necrosis factor receptor superfamily. Genomics. 62, 103–107 (1999).

48. Pispa, J., Mikkola, M. L., Mustonen, T. & Thesleff, I. Ectodysplasin, Edar and TNFRSF19 are expressed in complementary and overlapping patterns during mouse embryogenesis. Gene Expr. Patterns. 3, 675–679 (2003).

49. Chan, E. K., Nagaraj, S. H. & Reverter, A. The evolution of tropical adaptation comparing taurine and zebu cattle. Anim. Genet. 41, 467–477 (2010).

50. Hudson, N. J., Porto-Neto, L. R., Kijas, J., McWilliam, S., Taft, R. J. & Reverter, A. Information compression exploits patterns of genome composition to discriminate populations and highlight regions of evolutionary interest. BMC Bioinformatics. 15, 66 (2014).

51. Aftab, S., Semenec, L., Chu, J. S. & Chen, N. Identification and characterization of novel human tissue-specific RFX transcription factors. BMC Evol. Biol. 8, 226 (2008).

52. Blackshear, P. J., Graves, J. P., Stumpo, D. J., Cobos, I., Rubenstein, J. L. & Zeldin, D. C. Graded phenotypic response to partial and complete deficiency of a brain-specific transcript variant of the winged helix transcription factor RFX4. Development. 130, 4539–4552 (2003).

53. Fortes, M. R., et al. Gene network analyses of first service conception in Brangus heifers: use of genome and trait associations, hypothalamic-transcriptome information, and transcription factors. J. Anim. Sci. 90, 2894–2906 (2012).

54. Picardo, M. & Cardinali, G. The genetic determination of skin pigmentation: KITLG and the KITLG/c-Kit pathway as key players in the onset of human familial pigmentary diseases. J. Invest. Dermatol. 131, 1182–1185 (2011).

55. Gage, P. J., Qian, M., Wu, D. & Rosenberg, K. I. The canonical Wnt signaling antagonist DKK2 is an essential effector of PITX2 function during normal eye development. Dev. Biol. 317, 310–324 (2008).

56. Pausch, H., et al. Identification of QTL for UV-protective eye area pigmentation in cattle by progeny phenotyping and genome-wide association analysis. PLoS One. 7, e36346 (2012).

57. Sorbolini, S., et al. Use of canonical discriminant analysis to study signatures of selection in cattle. Genet. Sel. Evol. 48, 58 (2016).

58. Seo, K., Mohanty, T. R., Choi, T. & Hwang, I. Biology of epidermal and hair pigmentation in cattle: a mini-review. Vet. Dermatol. 18, 392–400 (2007).

59. Hagen, G., Müller, S., Beato, M. & Suske, G. Cloning by recognition site screening of two novel GT box binding proteins: a family of Sp1 related genes. Nucleic Acids Res. 20, 5519–5525 (1992).

60. Utsunomiya, Y. T., et al. Genome-wide mapping of loci explaining variance in scrotal circumference in nellore cattle. PLoS One. 9, e88561 (2014).

61. Kim, J., et al. The genome landscape of indigenous African cattle. Genome Biol. 18, 34 (2017).

62. Matukumalli, L. K., et al. Development and characterization of a high density SNP genotyping assay for cattle. PLoS One. 4, e5350 (2009).

63. Purcell, S., et al. PLINK: a tool set for whole-genome association and population-based linkage analyses. Am. J. Hum. Genet. 81, 559–575 (2007).

64. Decker, J. E., et al. Resolving the evolution of extant and extinct ruminants with high-throughput phylogenomics. Proc. Natl. Acad. Sci. U.S.A. 106, 18644–18649 (2009).

65. Zheng, X., Levine, D., Shen, J., Gogarten, S. M., Laurie, C. & Weir, B. S. A high-performance computing toolset for relatedness and principal component analysis of SNP data. Bioinformatics. 28, 3326–3328 (2012).

66. Alexander, D. H., Novembre, J. & Lange, K. Fast model-based estimation of ancestry in unrelated individuals. Genome Res. 19, 1655–1664 (2009).

67. Rosenberg, N. A. DISTRUCT: a program for the graphical display of population structure. Mol. Ecol. Notes. 4, 137–138 (2004).

68. Huson, D. H. & Bryant, D. Application of phylogenetic networks in evolutionary studies. Mol. Biol. Evol. 23, 254–267 (2006).

69. Patterson, N., et al. Ancient admixture in human history. Genetics. 192, 1065–1093 (2012).

70. Librado, P. & Rozas, J. DnaSP v5: a software for comprehensive analysis of DNA polymorphism data. Bioinformatics. 25, 1451–1452 (2009).

71. Bonfiglio, S., De Gaetano, A., Tesfaye, K., Grugni, V., Semino, O. & Ferretti, L. A novel USP9Y polymorphism allowing a rapid and unambiguous classification of Bos taurus Y chromosomes into haplogroups. Anim. Genet. 43, 611–613 (2012).

72. Edwards, C. J., et al. Dual origins of dairy cattle farming--evidence from a comprehensive survey of European Y-chromosomal variation. PLoS One. 6, e15922 (2011).

73. Günther, T. & Coop, G. Robust identification of local adaptation from allele frequencies. Genetics. 195, 205–220 (2013).

